# Centromeres in the thermotolerant yeast *K. marxianus* mediate attachment to a single microtubule

**DOI:** 10.1101/2025.01.24.634737

**Authors:** Daniel J. Barrero, Sabrine Hedouin, Yizi Mao, Charles L. Asbury, Andrew Stergachis, Eileen O’Toole, Sue Biggins

## Abstract

Eukaryotic chromosome segregation requires spindle microtubules to attach to chromosomes through kinetochores. The chromosomal locus that mediates kinetochore assembly is the centromere and is epigenetically specified in most organisms by a centromeric histone H3 variant called CENP-A. An exception to this is budding yeast which have short, sequenced-defined point centromeres. In *S. cerevisiae*, a single CENP-A nucleosome is formed at the centromere and is sufficient for kinetochore assembly. The thermophilic budding yeast *Kluyveromyces marxianus* also has a point centromere but its length is nearly double the *S. cerevisiae* centromere and the number of centromeric nucleosomes and kinetochore attachment sites is unknown. Purification of native kinetochores from *K. marxianus* yielded a mixed population, with one subpopulation that appeared to consist of doublets, making it unclear whether *K. marxianus* shares the same attachment architecture as *S. cerevisiae.* Here, we demonstrate that though the doublet kinetochores have a functional impact on kinetochore strength, kinetochore localization throughout the cell cycle appears conserved between these two yeasts. In addition, whole spindle electron tomography demonstrates that a single microtubule binds to each chromosome. Single-molecule nucleosome mapping analysis suggests the presence of a single centromeric nucleosome. Taken together, we propose that the *K. marxianus* point centromere assembles a single centromeric nucleosome that mediates attachment to one microtubule.

## Introduction

Organisms must accurately replicate and partition their genetic material during cell division. Eukaryotes achieve this using a megadalton protein machine called the kinetochore, which assembles on chromosomes and attaches to microtubules to segregate chromosomes during mitosis (Biggins, 2013; McAinsh and Marston, 2022). Chromosome segregation errors are a hallmark of cancers and many developmental diseases (Hanahan and Weinberg, 2011; Klaasen and Kops, 2022; Pfau and Amon, 2012), so it is critical to understand the underlying mechanisms that ensure accuracy.

Kinetochores assemble on chromosomal loci called centromeres (Mellone and Fachinetti, 2021; Talbert and Henikoff, 2020). Most centromeres are composed of kilobases to megabases of repetitive AT-rich DNA that lack sequence specificity and are epigenetically determined by CENP-A, a centromeric histone variant that replaces H3 in nucleosomes (Ali-Ahmad and Sekulic, 2020; Stirpe and Heun, 2023). In addition, most kinetochores bind to multiple microtubules. In contrast, point centromeres are sequence-based and best characterized in the budding yeast *Saccharomyces cerevisiae* where a 125-bp DNA sequence forms a single CENP-A (Cse4 in yeast) centromeric nucleosome that is sufficient to mediate kinetochore assembly (Furuyama and Biggins, 2007; Meluh et al., 1998; Palmer et al., 1987; Stoler et al., 1995). The budding yeast centromere has three <u>c</u>entromere <u>d</u>efining <u>e</u>lements (CDEI-III). While CDEI and CDEIII are short sequence-specific binding sites for the Cbf1 protein and the CBF3 complex, respectively, the CDEII element is a 79-88 base pair stretch of AT-rich DNA where Cse4 is positioned. Each *S. cerevisiae* kinetochore binds to a single microtubule emanating from the spindle pole body (SPB), the yeast centrosome equivalent (Kilmartin, 2014; Winey et al., 1995). Despite these differences, kinetochore components are structurally well conserved between yeast and humans, and it has been proposed that regional kinetochores are composed of repeat units of the kinetochores of point centromeres (Dendooven et al., 2023; Hamilton et al., 2019; Pesenti et al., 2022; Wigge and Kilmartin, 2001).

Budding yeast are a powerful model system to study kinetochore biology due to their relative simplicity. Native kinetochore particles can be readily purified from *S. cerevisiae* and have proven valuable for biophysical and biochemical analyses (Akiyoshi et al., 2010; de Regt et al., 2021; Miller et al., 2016). However, it has been difficult to perform structural biology on the purified kinetochore material due to its tendency to fall apart on electron microscopy grids (Gonen et al., 2012). To overcome this, we purified native kinetochore particles from the thermotolerant budding yeast *Kluyveromyces marxianus* as the adaptations evolved to live at high temperature tend to create more stable proteins that can be used for structural methods (Amlacher et al., 2011; Barrero et al., 2024; Meruelo et al., 2012; Szilágyi and Závodsky, 2000). The centromeres in *K. marxianus* contain the same centromere determining elements and CDE binding proteins as *S. cerevisiae*. However, the CDEII region of the *K. marxianus* centromere is 165 bp, double the length of CDEII in *S. cerevisiae* (Iborra and Ball, 1994). The functional reason for a longer CDEII element is unclear, but longer CDEIIs have evolved in multiple species of budding yeast and preserving the kinetochore interface appears to be the factor that allows these changes to be tolerated (Helsen et al., 2025). The longer CDEII element may also allow two centromeric nucleosomes to form. Consistent with this possibility, the kinetochores purified from *K. marxianus* appeared as two populations, a mixture of singlets and doublets. Treatment with nuclease reduced the doublet population, suggesting they are tethered by DNA (Barrero et al., 2024). A doublet kinetochore may have two separate microtubule binding sites or a single binding site containing additional microtubule binding complexes. An alternative explanation is that the doublet kinetochores are sisters that have maintained their connection through the purification. Distinguishing between these possibilities is not possible because the number of centromeric nucleosomes and microtubule binding sites that each *K. marxianus* kinetochore contains is not known.

Here, we addressed the structural properties of the *K. marxianus* kinetochore and mitotic spindle. We found that the spindle morphology and kinetochore positioning are similar to *S. cerevisiae*. Spindle reconstructions revealed too few microtubules per spindle pole to allow for multiple microtubules per centromere. Consistent with this, analysis of the chromatin accessibility landscape at centromeres using single molecule chromatin analysis confirmed the presence of a protected DNA region encompassing the entire centromere sequence, suggesting the presence of a single nucleosome. Taken together, our data suggest that each *K. marxianus* chromosome has a single centromeric nucleosome that mediates attachment to one microtubule.

## Results and Discussion

### Kinetochore doublets impact microtubule attachment strength

Earlier work demonstrated the presence of doublet and singlet kinetochores in purifications from *K. marxianus*, and that nuclease treatment reduced the proportion of doublets from approximately 40% to 20% (Barrero et al., 2024)(Figure 1A). To determine if the presence of doublets had a functional impact on the strength of kinetochores, we tested whether benzonase-treated kinetochores behave differently from untreated controls in a previously established optical trapping assay that assesses the functional strength of kinetochores (Akiyoshi et al., 2010; Barrero et al., 2024). We purified kinetochore particles from *K. marxianus* using a strain in which the Dsn1 kinetochore protein was tagged at the endogenous locus with 6xHis and 3xM3DK (Dsn1-His-M3DK) (Barrero et al., 2024). Logarithmically growing cells were treated with benomyl to suppress microtubule dynamics and enrich for mitotic cells, reducing cell cycle stage variability. Cell lysates were prepared and α-M3DK beads were added in the presence or absence of benzonase for 3 hours, after which the beads were thoroughly washed, and the kinetochores were eluted with M3DK peptide. Benzonase treatment did not dramatically alter kinetochore composition as assayed by silver stained SDS-PAGE, which demonstrated the retention of almost all protein bands in the benzonase-treated sample except bands above 150 kD (Figure 1B). We further analyzed kinetochore composition by mass spectrometry and confirmed members of every kinetochore subcomplex were present (Figure 1C, <u>MassIVE</u>). Thus, although the proportion of doublets is reduced with benzonase treatment, the overall protein composition of the kinetochores remained consistent.

**Figure 1.**
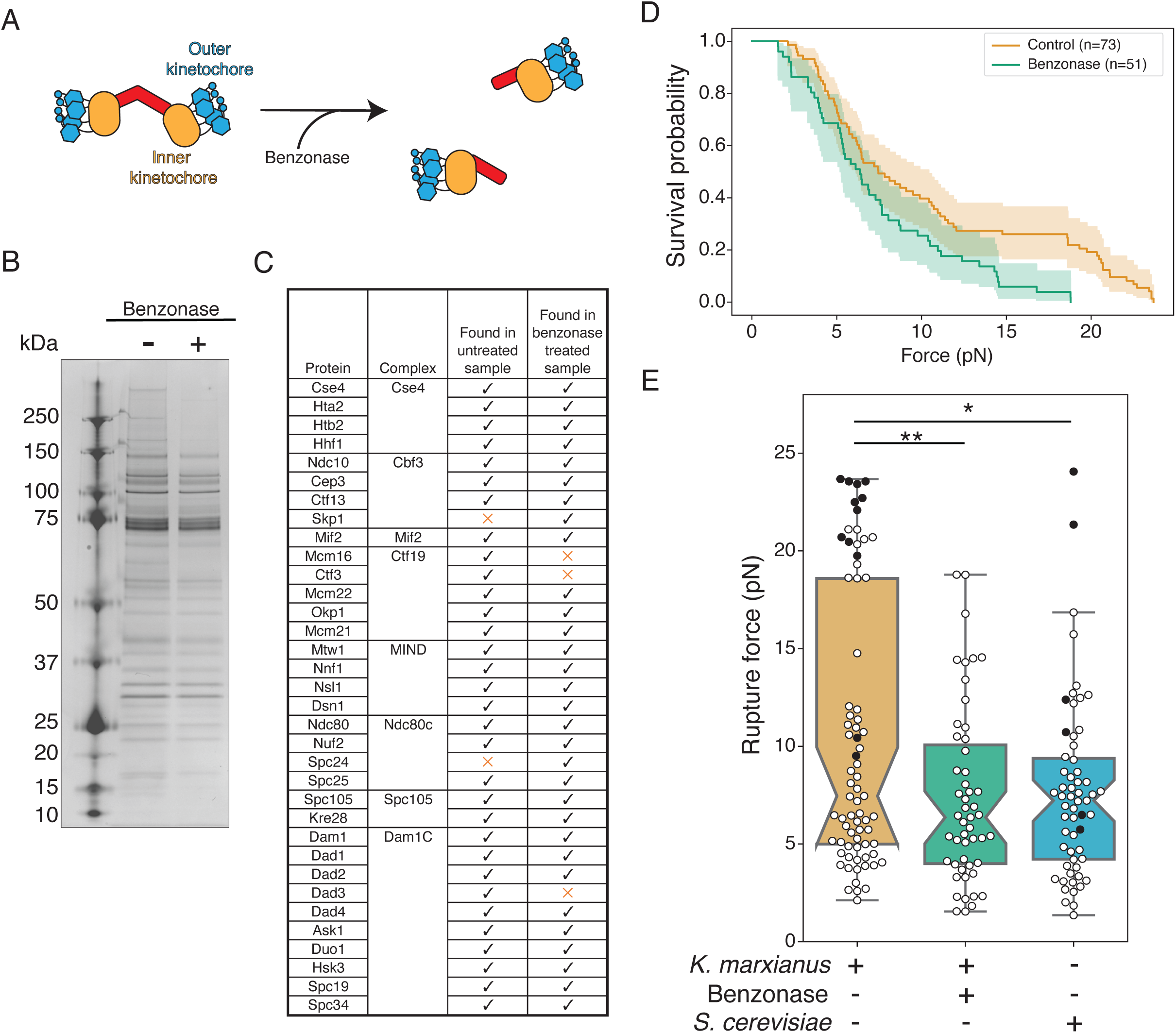
Benzonase treatment of purified kinetochores does not significantly impact kinetochore composition but affects kinetochore-microtubule attachment strength. A) Cartoons depict a proportion of purified *K. marxianus* kinetochores that appear as doublets (left) that can be converted to singlets (right) by benzonase nuclease treatment. B) Kinetochores purified via Dsn1-6His-3M3DK (from strain SBY18150) were visualized by silver-stained PAGE with or without benzonase treatment. C) Kinetochore proteins detected in a representative mass spectrometry analysis of purified kinetochores with or without benzonase treatment. Checks indicate the protein was detected, orange Xs indicate it was not. D) Survival probability curves of force ramp experiments of untreated *K. marxianus* kinetochores (orange, median = 7.5 pN) and benzonase treated *K. marxianus* kinetochores (green. median = 6.4 pN). Shaded regions represent the 95% confidence intervals. The survival curves differ significantly (p < 0.005 by log-rank test). E) Box plots of rupture force values for kinetochore-microtubule attachments overlayed with individual events for untreated *K. marxianus* (orange), benzonase treated *K. marxianus* (green), and untreated *S. cerevisiae* (blue, median = 7.2). White circles represent true ruptures, and black circles represent escape events. Untreated and benzonase treated *K. marxianus* differed significantly (p<0.005, log-rank test), as did untreated *K. marxianus* and untreated *S. cerevisiae* (p=0.04, log-rank test). Benzonase treated *K. marxianus* and untreated *S. cerevisiae* were indistinguishable (p=0.77, log-rank test).

We next performed optical trapping on the kinetochores treated with and without benzonase. Despite their similar composition, benzonase-treated kinetochores were significantly weaker than untreated controls (Figure 1D, E, p < 0.005 by log-rank test). The benzonase-treated kinetochores also lost the bimodal distribution present in untreated controls and their rupture force was indistinguishable from *S. cerevisiae* kinetochores (Figure 1E, p = 0.77 by log-rank test), suggesting the stronger kinetochores are doublets. These results are consistent with the possibility of multiple microtubule binding units in a doublet kinetochore.

### Kinetochore positioning throughout mitosis is conserved between K. marxianus and S. cerevisiae

Given the possibility that the *K. marxianus* kinetochore-microtubule attachment site is different from *S. cerevisiae,* we sought to determine if the overall spindle morphology and kinetochore positioning were conserved. In *S. cerevisiae*, kinetochores remain assembled and attached to microtubules throughout the cell cycle except for a short time during S phase (Guacci et al., 1997; Jin et al., 2000; Kitamura et al., 2007; Tanaka et al., 2002; Winey and O’Toole, 2001). This causes kinetochores to cluster near the SPB for most of the cell cycle. Because the kinetochores are clustered, they are not individually resolvable and appear as a single focus prior to spindle formation. Once the SPB duplicates and separates to form short spindles, the kinetochores cluster into two foci. We asked whether *K. marxianus* kinetochores behave similarly by epitope tagging the outer kinetochore protein Dad1 at the endogenous locus with the fluorophore mKate2 (Dad1-mKate2) and performing microscopy. Cells were visible in all stages of budding, indicating active progression through the cell cycle (Figure 2A). As in *S. cerevisiae*, unbudded cells showed a single focus of Dad1 fluorescence that was localized to a single mass of DNA (Figure 2A, unbudded). Small-budded cells maintained a single kinetochore focus and amorphous nuclear DNA signal (Figure 2A, small budded). Occasionally, small-budded cells could be seen with two kinetochore foci, indicating spindle formation and entry into mitosis. In large-budded cells, two kinetochore puncta were clearly visible. Large-budded cells were categorized as pre-anaphase if their DNA was rounded and still in the mother cell, and anaphase if it was elongated or separated (Figure 2A, large-budded/pre-anaphase, large-budded/anaphase). Pre-anaphase inter-kinetochore distance measurements had a median distance of 0.420 µm, which is shorter than the 0.6 µm distance in *S. cerevisiae* (Figure 2B)(Winey and O’Toole, 2001).

**Figure 2.**
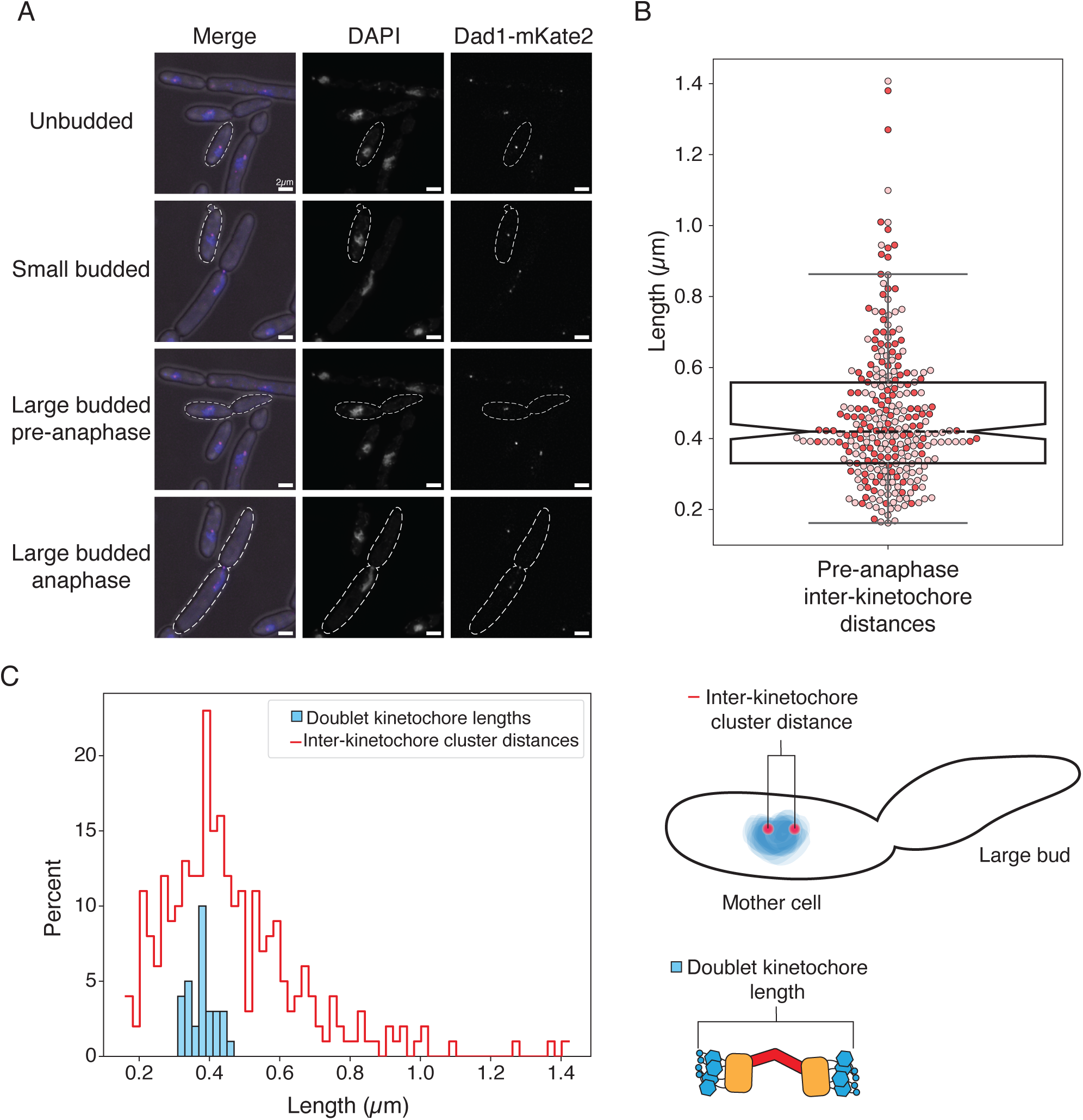
Characterization of *K. marxianus* kinetochore positioning *in vivo*. A) Representative images of DAPI-stained Dad1-mKate2 *K. marxianus* cells throughout the cell cycle. Kinetochores cluster into one or two foci. Scale bar is 2 µm. B) Pre-anaphase inter-kinetochore cluster distances measured as the distance from the center of one Dad1-mKate2 focus to the center of the second (median = 0.42 µm, n = 274 distances from 2 biological replicates). Each circle is a single measurement, colored by biological replicate, and the boxplot notch represents the median value. C) Left: A histogram comparison of the inter-kinetochore cluster distance measured by fluorescence and the doublet kinetochore lengths previously measured by tomography (Barrero et al., 2024), indicating that these two measurements are consistent. Right: cartoon schematics of the inter-kinetochore cluster distance measurement (top) and the doublet kinetochore length measurement.

One possibility for the nature of the doublet kinetochores is that they represent sisters. In this scenario, the inter-kinetochore cluster distance should accommodate the measurements of doublet kinetochores previously obtained by cryo-electron tomographic (cryo-ET)(Barrero et al., 2024). We therefore compared the distribution of end-to-end length measurements of doublet kinetochores from previously published cryo-electron tomography (cryo-ET) data to the distribution of inter-kinetochore distances measured by light microscopy (Figure 2C)(Barrero et al., 2024). The cryo-ET doublet kinetochore lengths (375 nm, n=31 doublet kinetochores) easily fit within the spread of inter-kinetochore cluster distances, and cluster within 100 nm of the median inter-kinetochore distance (Figure 2C, 420 nm, n=274 measurements). Thus, kinetochore positioning through mitosis is consistent between *S. cerevisiae* and *K. marxianus*, and previously published doublet kinetochore lengths can be easily accommodated within the inter-kinetochore cluster distances.

### Spindle reconstructions indicate a single microtubule per centromere

To determine the number of microtubule binding sites at the *K. marxianus* kinetochore, we reconstructed whole mitotic spindles from logarithmically growing *K. marxianus* cells. As shown previously, electron tomography provides a detailed view of spindle morphology in yeast cells because it allows for the clear resolution of individual microtubules and spindle poles (O’Toole et al., 2017a; O’Toole et al., 2017b; O’Toole et al., 1999). We therefore prepared cells by <u>H</u>igh <u>P</u>ressure <u>F</u>reezing and <u>F</u>reeze <u>S</u>ubstitution (HPF/FS) and collected electron tomograms to reconstruct their spindles.

We initially focused on the SPB, which we expected to appear similarly to that of *S. cerevisiae*, as a tri-laminar disk with a heavy staining central plaque embedded within the nuclear membrane and an inner and outer plaque from which nuclear and cytoplasmic microtubules emanate (Byers and Goetsch, 1975; O’Toole et al., 1997; O’Toole et al., 1999). Indeed, structures resembling the trilaminar *S. cerevisiae* SPB were readily identifiable in the nuclear membrane of dividing cells (Figure 3A). The central layer, commonly referred to as the central plaque, is embedded in the nuclear envelope and has a layered structure like *S. cerevisiae* (Figure 3A, yellow arrows). However, the SPB is consistently smaller in size with an average diameter of 66 nm compared to roughly 100 nm in *S. cerevisiae* (Byers and Goetsch, 1974). Mother SPBs in *K. marxianus* also seemed to assemble a tilted outer plaque on the cytoplasmic face (Figure 3A, blue arrows) and half bridges that are the location of new SPB assembly were visible (Figure 3A, green arrows)(O’Toole et al., 1999).

**Figure 3.**
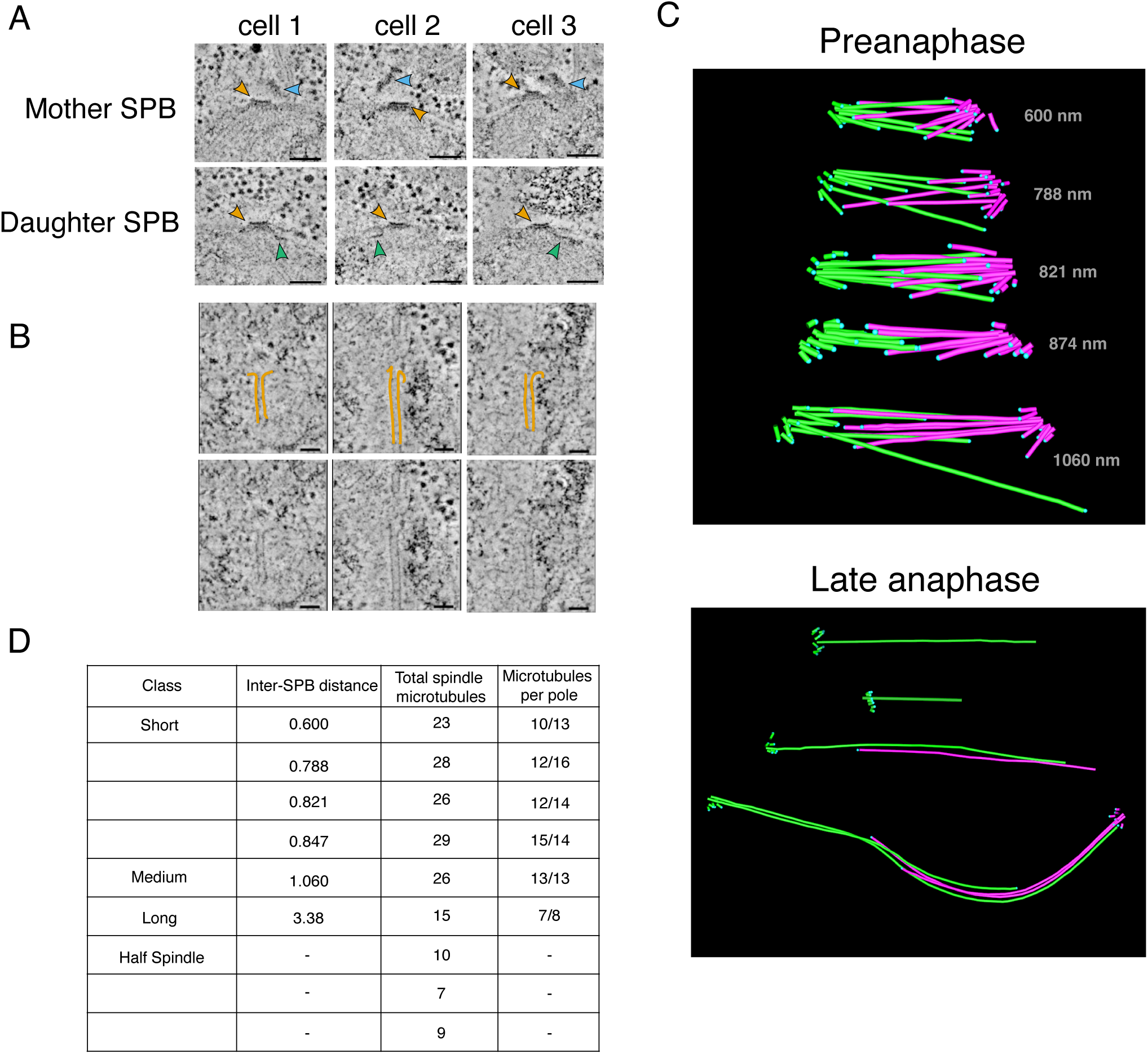
3-dimensional reconstructions of *K. marxianus* mitotic spindles are consistent with a single microtubule binding site per kinetochore. A) An array of mother (top row) and daughter (bottom row) spindle pole bodies from three *K. marxianus* cells. Yellow arrows indicate the spindle pole bodies, blue arrows indicate the tilted outer plaques of mother SPBs. Half bridges are visible as a thin line of dark staining extending along the nuclear membrane (green arrows). Scale bars represent 100 nanometers. B) Representative gallery of spindle microtubule tips. Protofilaments have been traced in yellow (top row), and unmarked images have been included for comparison (bottom row). Scale bars represent 50 nanometers. C) Examples of reconstructed spindle models. Preanaphase spindles are shown on the top and late anaphase half spindles, and a complete late anaphase spindle is shown on the bottom. Microtubules originating from the same SPB are denoted in the same color, either pink or green. Length in nanometers from SPB to SPB are denoted on each preanaphase spindle. Blue dots indicate the termination point of the microtubule. The colored asterisks correspond to the spindles in the table in (D). D) Classes, lengths, and microtubule content of reconstructed mitotic spindles. As length was measured from SPB to SPB, this measurement was omitted for half spindles where only one SPB was visible.

To reconstruct the spindle, we focused on microtubules originating in the SPB and extending into the nucleus. For microtubules with discernable plus ends, we were able to make out limited tip morphology, including some examples of protofilament flares (Figure 3B). *K. marxianus* has 8 chromosomes as opposed to 16 in *S. cerevisiae*. Therefore, if there is only one microtubule attachment per chromosome like *S. cerevisiae*, we expected to see 8 kinetochore microtubules and roughly 3 to 8 interpolar microtubules per SPB based on *S. cerevisiae* spindle reconstructions (Winey et al., 1995). Nine mitotic spindles were reconstructed, measuring from 0.6 µm to 3.38 µm in length, defined as the SPB-to-SPB distance (Figure 3C). The four short spindles (defined as less than 1 µm long) and one medial (∼1 µm) spindle averaged 27 microtubules total, with individual SPBs containing between 10 and 15 microtubules (Figure 3D). In *S. cerevisiae*, short and medial spindles contain 38-43 spindle microtubules (Winey et al., 1995). Four of the spindles were in late anaphase, and for three of those we reconstructed only a half spindle. The complete late anaphase spindle contained 15 microtubules total, 8 from one spindle pole and 7 from the other (Figure 3D). This is in stark contrast to published data from *S. cerevisiae* where the anaphase B spindle contains ∼39 mts (Winey et al., 1995). The reduced number of microtubules compared to *S. cerevisiae* is consistent with the smaller SPBs, as SPB surface area tends to scale with number of microtubules (Storchova et al., 2006). Considering *K. marxianus* has 8 chromosomes, these spindle reconstructions are most consistent with roughly one microtubule per kinetochore and a few additional microtubules that serve as interpolar microtubules.

### K. marxianus appears to have a single centromeric nucleosome

Our finding that there is only a single microtubule that binds to each kinetochore suggested that the doublet kinetochores might represent sisters or two kinetochore units that simultaneously interact with a single microtubule. Tomographic reconstructions in yeast are unable to distinguish this as the chromosomes and kinetochores do not condense into a visible structure that can be readily identified in the EM. To address this, we sought to establish the number of centromeric nucleosomes by single-molecule chromatin fiber sequencing (Fiber-seq), which can map nucleosome location on DNA with high resolution by detecting m6A DNA methylation on open chromatin through long read sequencing (Stergachis et al., 2020). Briefly, logarithmically growing cells are spheroplasted and treated with a N6-adenine DNA methyltransferase (m6A-MTase) which nonspecifically methylates accessible DNA. Individual m6A-labeled chromatin fibers are sequenced using long-read single-molecule sequencing to >1000x genomic coverage, which enables the identification of single-molecule protein occupancy events at near single-nucleotide resolution (Figure 4A). Since sequenced reads are >10kb in length, the entirety of all eight *K. marxianus* point centromeres was readily captured along a sequenced fiber. If *K. marxianus* assembles a single centromeric nucleosome we would expect an unmethylated protected DNA region encompassing the entire centromere whereas if two centromeric nucleosomes were present, we would expect two protected regions in close proximity, separated by methylated linker DNA (Figure 4A).

*K. marxianus* single-molecule chromatin fiber reads revealed well-protected regions that mapped to the whole length of all 8 centromeres and were almost devoid of m6A-MTase methylation, as observed for *S. cerevisiae* (Figure 4B). These regions showed extremely high nucleosome occupancy which peaks at the center of the centromere sequence (Figure 4C). The centromere nucleosome footprint for *K. marxianus* was larger than that of *S. cerevisae* (median of 227 bp versus 165 bp for *K. marxianus* and *S. cerevisiae*, respectively), likely due to its extended CDEII region, however this protected region is still only large enough to account for a single centromeric nucleosome, albeit with a potentially different amount of DNA wrapping due to its longer CDEII (Figure 4D) (Iborra and Ball, 1994). This data confirms the presence of a single nucleosomal footprint at the centromere, and taken with the tomographic reconstructions suggests a point centromere organization similar to *S. cerevisae*.

**Figure 4.**
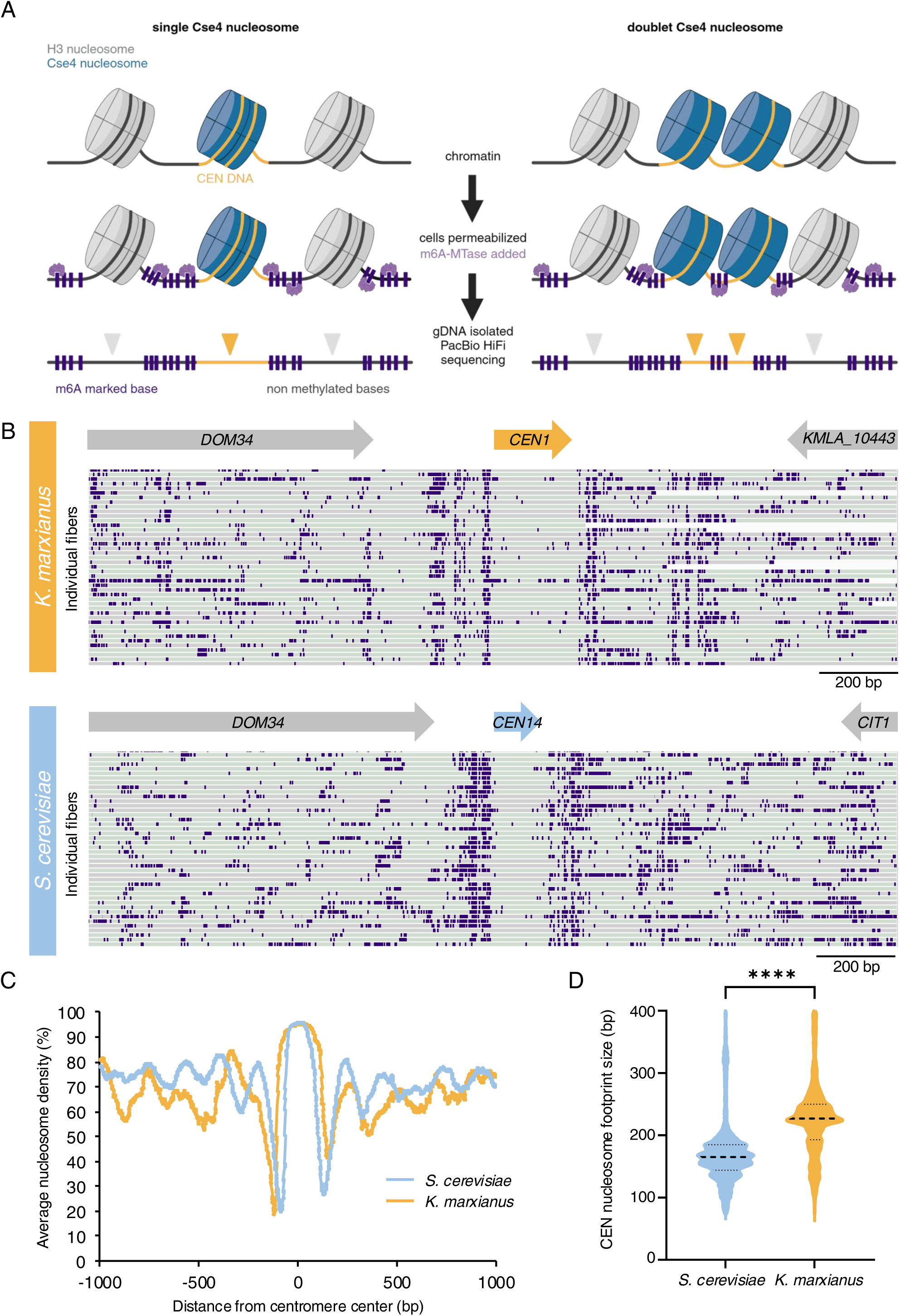
Fiber-Seq analysis confirms a single centromeric nucleosomal footprint. A) A schematic of the Fiber-seq protocol, in which cells are permeabilized and treated with Hia5 m6A-methyltransferase before having their gDNA purified and sequenced by PacBio HiFi sequencing to reveal a high-resolution map of DNA protection via the presence or absence of methylated DNA bases. If there is only one centromeric nucleosome, we expect the centromere DNA to remain fully unmethylated (left panel), whereas, if two distinct nucleosomes are present at the centromere, we expect to detect methylated within the centromere corresponding to the linker DNA (right panel). B) Examples of individual Fiber-seq reads through the centromere of chromosome I mapped onto a *K. marxianus* reference genome (DMKU3-1042). Individual reads (gray and green bars depending on read strand orientation) are stacked vertically, and CEN 1 location is marked (upper panel). Examples of individual Fiber-seq reads through the centromere of the syntenic region on chromosome XIV in *S. cerevisiae* are shown on the lower panel. Purple boxes indicate methylation signal. C) A comparison of the average nucleosome occupancy at all CENs in *K. marxianus* (yellow line) and *S. cerevisiae* (blue line). D) A comparison of the size of the nucleosome footprint overlapping with the centromere in *S. cerevisiae* and *K. marxianus*. The black dotted lines represent the median length (165 bp for *S. cerevisiae* and 227 bp for *K. marxianus*), and lower and upper quartiles from ∼11,000 fibers analyzed. **** indicates significant difference between the two species with unpaired two-tailed t-test (P-value <0.0001).

### Concluding remarks and perspectives

Here we report the first ultrastructural analysis of spindle organization of *K. marxianus*. Because the CDEII element of the *K. marxianus* centromere is longer than *S. cerevisiae*, it was unclear whether multiple centromeric nucleosomes and microtubule binding sites exist. In addition, purification of native *K. marxianus* kinetochores contained “doublets” that impacted kinetochore strength, consistent with both kinetochore units interacting with microtubules as in a regional centromere (Barrero et al., 2024). To determine if the mitotic spindle is different from *S. cerevisiae*, we examined kinetochore localization and cell morphology throughout the *K. marxianus* cell cycle to identify potential differences. However, we found that kinetochore localization throughout the cell cycle was similar to *S. cerevisiae*. Consistent with this, we reconstructed whole mitotic spindles from electron tomograms, and they revealed an insufficient number of microtubules per chromosome for regional centromeres. Additionally, Fiber-seq analysis examining chromatin accessibility at high resolution determined there is a single nucleosomal footprint at the *K. marxianus* centromere. While it is possible that there are two stacked Cse4 octamers or a higher proportion of wrapped DNA, our data is most compatible with a single nucleosome, which is consistent with a recent study suggesting that longer yeast centromeres evolve as long as they can maintain kinetochore interface interactions (Helsen et al., 2025). Thus, despite the appearance of doublet kinetochores, *K. marxianus* centromeres have a single microtubule binding site. The role and origin of the doublet kinetochores is therefore still unclear. Although it is possible that a subset of sister kinetochores remained linked throughout our kinetochore purification process, we did not detect cohesin, the complex responsible for linking sister chromatids during mitosis, in the mass spectroscopy analysis of these kinetochore purifications (Barrero et al., 2024). It will be important to determine if the doublets represent a physiological unit of kinetochores in the future. It will also be interesting to understand the expanded CDEII of *K. marxianus* and its implications for nucleosome wrapping and kinetochore assembly.

## Materials and methods

### Strain construction

The *Saccharomyces cerevisiae* strain used in this study is SBY8253 (*DSN1-6xHis-3xM3DK:URA3*) and was derived from the W303 background and was previously described (Barrero et al., 2024). The *Kluyveromyces marxianus* strains used in this study are SBY18150 (*DSN1-6xHis-3xM3DK:KanMX*) and SBY22682 (*DAD1-mKATE2:KanMX*) which were derived from SBY17411 (NRRL Y-8281, USDA ARS culture collection). All strains were tagged at the endogenous locus via integration. Briefly, DNA fragments of either 500 or 1000 bases immediately upstream and downstream of the desired integration site were generated from genomic DNA. A backbone plasmid was selected based on the desired tags, and the fragments amplified from genomic DNA were inserted into the backbone plasmid via Gibson assembly such that each plasmid contained a restriction site, followed by 500 to 1000 base pairs of upstream homology, followed by the desired tags and markers, followed by 500 to 1000 base pairs of downstream homology, followed by another restriction site. Plasmids were then digested and transformed into the desired strain for integration by homologous recombination. Successful integration was confirmed by PCR and immunoblotting. The plasmids (pSB prefix) and yeast strains (SBY prefix) used are as follows: SBY18150 contains plasmid pSB2951 generated with primers SB5736, SB5737, SB5738, SB5739, transformed into SBY17411. SBY22682 contains plasmid pSB3501 generated with primers SB8274, SB8275, SB8276, SB8277, SB8278, SB8279, transformed into SBY17411. All tagged strains grew similarly to the parent strain.

### Yeast growth and kinetochore purification for optical trapping

All yeast growth was performed as described previously (Barrero et al., 2024). Briefly, yeast were grown in YPD (1% yeast extract, 2% peptone, 2% D-glucose). SBY18150 cultures were grown in the presence of 200 µg/ml G418. Large cultures were grown on shakers (220 rpm) at 22 °C or 30 °C for *S. cerevisiae* and *K. marxianus,* respectively. Cultures were treated with benomyl at a final concentration of 30 µg/ml (1:1 addition of 60 µg/ml benomyl YEP media) for 2 hours at 23 °C and then harvested by centrifugation for 10 minutes at 5000xg at 4 °C. Kinetochores were purified as previously described (Barrero et al., 2024). Briefly, the endogenous *DSN1* kinetochore gene was C-terminally tagged with 6xHis and 3xM3DK. Harvested yeast were resuspended in Buffer H (25 mM HEPES pH 8.0, 150 mM KCl, 2 mM MgCl2, 0.1 mM EDTA pH 8.0, 0.1% NP-40, 15% glycerol) supplemented with protease inhibitors, phosphatase inhibitors, and 2 mM DTT. After resuspension and re-spinning, yeast pellets were frozen in liquid nitrogen and lysed using a Freezer Mill (SPEX, Metuchen NJ). Lysate was clarified via ultracentrifugation at 24,000 RPM (98,000 x g) for 90 minutes and the protein layer was extracted with a syringe. This extract was incubated with magnetic α-M3DK antibody conjugated Dynabeads (Invitrogen, Waltham MA) for 3 hours at 4 °C with rotation. For benzonase treated samples, 300 units of benzonase per milliliter of lysate were added during this incubation step. Dynabeads were washed with 10x bead volume of Buffer H 5 times (the last 3 washes omitting DTT and phosphatase inhibitors). Kinetochores were eluted with 0.5 mg/ml 3xM3DK peptide in Buffer H lacking DTT and phosphatase inhibitors. For mass spectrometry, kinetochores were eluted from Dynabeads with 0.2% RapiGest (Waters Corporation, Milford MA) in 50 mM HEPES pH 8.0. For all experiments, the total protein concentration was determined by NanoDrop measurement and the relative purity by silver stain gel analysis.

### Optical trapping

Optical trapping rupture force assays were performed as previously described (Barrero et al., 2024). Streptavidin coated 440 nm polystyrene beads (Spherotech, Lake Forest IL) were functionalized with biotinylated α-penta-His antibody (Qiagen, Hilden Germany or R&D Systems, Minneapolis MN) and stored in BRB80 containing 8 mg/ml BSA and 1 mM DTT at 4 °C with continuous rotation. Beads were decorated with purified kinetochores (via Dsn1-6His-3M3DK) in a total volume of 20 μl incubation buffer (BRB80 containing 1.5 mg/mL κ-casein). To ensure sparse decoration of the beads and reduce the likelihood of multiple kinetochore-microtubule interactions being measured simultaneously, we empirically determined kinetochore concentrations such that roughly 1 in 10 beads exhibited microtubule binding activity during the assay. Dynamic microtubule extensions were grown from coverslip-anchored GMPCPP-stabilized microtubule seeds in a microtubule growth buffer consisting of BRB80, 1 mM GTP, 250 µg/ml glucose oxidase, 25 mM glucose, 30 µg/mL catalase, 1 mM DTT, 1.4-1.5 mg/mL purified bovine brain tubulin and 1 mg/mL κ-casein. Assays were performed at 23 °C. Rupture force experiments were performed as in (Barrero et al., 2024). Briefly, an optical trap was used to apply a force of ∼1-2 pN in the direction of microtubule assembly. Once beads were observed to track with microtubule growth for roughly 30 seconds (to ensure end-on attachment), the applied force was increased at a constant rate of 0.25 pN/s until bead detachment. Records of bead position over time were generated and analyzed using custom software (LabVIEW and Igor Pro, respectively) and used to determine the rupture force, which was marked as the maximum force sustained by the attachment during each event.

### Cell fixation and Microscopy

Briefly, SBY22682 was grown in YPD (1% yeast extract, 2% peptone, 2% D-glucose) at 22 °C. 1 ml was removed during mid-log phase growth (∼OD 0.6) and yeast were pelleted by centrifugation at 21,000xg for 1 minute. Supernatant was removed and the yeast pellet was resuspended in 1 ml 0.1 M potassium phosphate pH 6.4 with 3.7% formaldehyde for fixation. This mixture was incubated at room temperature for 10 min before centrifugation at 21,000xg for 1 minute. Supernatant was removed and fixed cells were resuspended in 1 ml 0.1 M potassium phosphate pH 6.4 and stored at 4 °C for up to two weeks. Immediately prior to imaging, cells were pelleted with the same spin parameters and resuspended in 100 µl DAPI staining buffer (1.2 M sorbitol, 1% Triton-X100, 0.1 M potassium phosphate pH 7.5, 2 µg/ml DAPI) for 10 minutes. Cells were then pelleted again and resuspended in 100 µl imaging buffer (1.2 M sorbitol, 1% Triton-X100, 0.1 M potassium phosphate pH 7.5). Cells were applied to microscope slides with thin agarose pads (Skinner et al., 2013).

Fixed cells were imaged using a Deltavision Ultra deconvolution high resolution microscope with a 100x/1.42 PlanSApo oil immersion objective (Olympus Life Science, Waltham, MA). Images were collected with a 16-bit sCMOS detector. Cells were imaged using Z-stacks with 0.2µm steps through the entire cell. Deconvolution was done using standard settings through SoftwoRX software. All quantification was done using ImageJ (National Institutes of Health).

### Tomography

Cells were prepared for electron tomography using high pressure freezing followed by freeze substitution as previously described (Giddings et al., 2001; O’Toole et al., 2017b; O’Toole et al., 1999). Briefly, logarithmically growing *K. marxianus* cells were collected by vacuum filtration and frozen using a Leica EMPact2 high pressure freezer (Leica Biosystems, Deer Park IL). The frozen cells were then freeze substituted in 1% OsO_4_ and 0.1% uranyl acetate in acetone and embedded in epon. Thick (250 nm) sections were collected onto formvar-coated slot grids. The grids were then stained with 2% uranyl acetate followed by Reynolds lead citrate and 15nm gold particles (BBI International) were affixed to the section surface to serve as fiducial markers for alignment.

Tomography was performed using a Tecnai F30 microscope operating at 300 kV (Thermo Fisher, Waltham MA). Dual-axis tilt series (+/-60°, imaged every 1.5°) were collected using SerialEM software (Mastronarde, 2005) and a Gatan OneView camera at a pixel size of 1.5 nm. For most data sets, tilt series were collected from 2-3 serial sections to reconstruct the entire mitotic spindle in the tomographic volume. Serial tomograms were computed and joined using the IMOD 4.9 software package (Kremer et al., 1996; Mastronarde, 1997; Mastronarde and Held, 2017).

Spindle microtubules from either pole were tracked and their plus ends modeled using the 3dmod program of the IMOD software package. The spindles were then projected in 3D to show their arrangement within the volume. Measurements of spindle length, spindle microtubule length and diameter of spindle pole bodies were collected using the imodinfo program in the IMOD software package. In total, 6 complete spindles (ranging in length from 0.6 mm to 3.38 mm) and 3 half spindles from late anaphase were reconstructed.

### Fiber-seq

Yeast Fiber-seq was performed as described (Popchock et al., 2023). Briefly, 10 mL of *K. marxianus* cells (SBY18150) were grown in YPD media to mid-log phase and harvested by centrifugation. Cells were washed once with cold dH2O and resuspended in cold KPO4/Sorbitol buffer (1 M Sorbitol, 50 mM Potassium phosphate pH 7.5, 5 mM EDTA pH 8.0) supplemented with 0.167% β-Mercaptoethanol. Cells were spheroplasted by addition of 0.15 ug/mL final concentration of Zymolyase T100 (Amsbio) and incubated at 23 °C for 15 min on a roller drum. Spheroplasts were pelleted at 1200 rpm for 8 min at 4 °C, washed twice with cold 1M Sorbitol, and resuspended in 58 µL of Buffer A (1M Sorbitol, 15 mM Tris-HCl pH 8.0, 15 mM NaCl, 60 mM KCl, 1 mM EDTA pH 8.0, 0.5 mM EGTA pH 8.0, 0.5 mM Spermidine, 0.075% IGEPAL CA-630). Spheroplasts were treated with 1 µL of Hia5 MTase (200U) and 1.5 µL of 32mM S-adenosylmethionine (NEB) for 10 min at 25 °C. Reaction was stopped by addition of 3 µL of 20% SDS (1% final concentration) and high molecular weight DNA was purified using the Promega Wizard® HMW DNA extraction kit (A2920).

HMW Fiber-seq modified gDNA was sheared using a g-TUBE (520079) and PacBio SMRTbell libraries were then constructed and sequenced as described (Bohaczuk et al., 2024). Circular consensus sequence reads were generated from raw PacBio subread files and processed as described (Vollger et al. 2024). Reads were mapped to the DMKU3-1042 *K. marxianus* reference genome (NCBI RefSeq assembly GCF_001417885.1). Nucleosomes were then defined using the default parameters of *fibertools* (PMID 38849157). Fibers overlapping with the center of each eight centromeres +/- 1000bp were extracted. The nucleosome density was calculated by counting the number of nucleosomes that overlap with each base pair of the region of interest divided by the number of fibers overlapping with that position. The nucleosome density of all eight centromeric regions were averaged and plotted in figure 4C. The size of the centromeric nucleosome footprint was extracted using *bedmap –echo-map-size*. Values superior to 400 bp were filtered out as they might result from incomplete m6A methylation. Statistical analysis was performed using Prism. Fiber-seq data from *S. cerevisiae* were taken from (Dubocanin et al., 2024; Popchock et al., 2023).

## Data availability

Mass spectrometry data generated in this study is being made available through <u>Mass</u> Spectrometry Interactive <u>V</u>irtual <u>E</u>nvironment (MassIVE, University of California San Diego, MSV000095817): https://massive.ucsd.edu/ProteoSAFe/dataset.jsp?task=39a41e7265684e05ad04070428284442. The *K. marxianus* sequencing data generated in this study have been deposited in the NCBI Sequence Read Archive (SRA) database under the accession number PRJNA1194201. *S. cerevisiae* Fiber-seq data are available at SRA under accession number PRJNA1189155 (Popchock et al., 2025).

## Acknowledgments

Electron microscopy was performed at the University of Colorado, Boulder EM Services Core Facility in the MCDB Department, with the technical assistance of facility staff. We thank Shane Neph for his assistance with the Fiber-seq analysis, Steve MacFarlane for help with tomography cell preparation and Christian Nelson for help with strain construction. This work was supported by NIH DP5-OD029630 to A.B.S. who also holds a Career Award for Medical Scientists from the Burroughs Wellcome Fund and is a Pew Biomedical Scholar, NIH R35GM134842 to CLA, and NIH R35 GM149357 to SB who is also an investigator of the Howard Hughes Medical Institute.

